# GTPase-activating protein ARAP1 regulates circular dorsal ruffles as a nutrient uptake mechanism in the Hep3B hepatocellular carcinoma cell line

**DOI:** 10.1101/2023.12.31.573800

**Authors:** Xiaowei Sun, Yanan Li, Yuxin He, Longjiao Cheng, Jinzi Wei, Linxuan Du, Zhongyang Shen, Sei Yoshida

## Abstract

Circular dorsal ruffles (CDRs), large-scale rounded membrane ruffles, function as precursors of macropinocytosis. We recently reported that CDRs are exposed in the Hep3B hepatocellular carcinoma cell line, while not in other hepatocellular carcinoma cell lines, indicating that the CDRs in Hep3B are associated with malignant potential. In this study, we investigated the cellular function of CDRs in Hep3B cells by focusing on the molecular mechanisms of the GTPase-activating protein ARAP1. ARAP1 was localized to the CDRs, the sizes of which were reduced by deletion of this protein. High-resolution scanning electron micrographs revealed that CDRs comprise small vertical lamellipodia, the expression pattern of which was disrupted in ARAP1 KO cells. Extracellular solute uptake, rate of cell growth, and malignant potential were attenuated in the KO cells. ARAP1 is also localized in Hep3B cell mitochondria, although not in those of the Huh7 hepatocellular carcinoma cell line. On the basis of these findings, we propose that the aberrant expression of ARAP1 in Hep3B cells modulates CDRs, thereby resulting in an excess uptake of nutrients as an initial event in cancer development.

**SUMMARY STATEMENT:** ARAP1 regulates circular dorsal ruffles (CDRs) in the Hep3B HCC cell line and deletion of this protein attenuates malignant potential, thereby indicating the involvement of CDRs in cancer development.

## INTRODUCTION

Circular dorsal ruffles (CDRs) are large-scale rounded membrane ruffles evoked on the dorsal surface of cells (Hoon et al., 2012; Itoh and Hasegawa, 2013; Yoshida et al., 2018b). In response to stimulation with extracellular ligands, such as platelet-derived growth factor (PDGF), epidermal growth factor (EGF), hepatocyte growth factor (HGF), and insulin; cells induce the surface development of vertical membrane ruffles to produce these CDRs, in which the ruffles bend inward and gradually close the open area to generate vesicles (Yoshida et al., 2018b). It has accordingly been proposed that CDRs are involved in macropinocytosis, a large-scale type of endocytosis that facilitates the uptake of extracellular solutes (Gu et al., 2011; Hoon et al., 2012; Itoh and Hasegawa, 2013). Live-cell imaging of CDRs in mouse embryonic fibroblasts (MEFs) and human glioblastoma LN299 cells clearly show that large-sized vesicles (macropinosomes) are generated upon the closure of CDRs (Yoshida et al., 2018b; Zdzalik-Bielecka et al., 2021). Since CDRs were originally identified in PDGF-stimulated human glial cells in 1983 (Mellstroom et al., 1983), the structures have been confirmed in more than 15 different cell types, including MEFs (Yoshida et al., 2018b), NIH-3T3 cells (Hasegawa et al., 2012), kidney epithelial cells podocytes (Hua et al., 2023), LN299 cells (Zdzalik-Bielecka et al., 2021), and the Hep3B hepatocellular carcinoma (HCC) cell line (Sun et al., 2022).

In terms of the molecular mechanism of CDR formation, numerous studies have tended to focus on the roles of small GTPases (Itoh and Hasegawa, 2013). For example, Rac1 has been detected in CDRs (Hooshmand-Rad et al., 1997), and it has been demonstrated that whereas overexpression of the active form of Rab5a induces CDRs, this effect was diminished by expression of the dominant-negative form of Rac (Palamidessi et al., 2008). Moreover, co-expression of the dominant-active form of Ras and wild-type Rab5a has been shown to induce CDRs in MEFs (Lanzetti et al., 2004). Similarly, the role of ARF small GTPases and associated signaling molecules have been demonstrated. To date, six different ARFs have been identified in mammalian cells, although in humans, who lack ARF2, only five of these are expressed (Sztul et al., 2019). Among these, ARF1, 5, and 6 have been shown to be localized in CDRs (Hasegawa et al., 2012; Zobel et al., 2018). In common with other small GTPases, ARFs are negatively regulated by GTPase-activating proteins (GAPs), of which more than 15 ARF GAPs have been reported, with some being found to be involved in CDR formation (Tanna et al., 2019). Overexpression studies have revealed that ACAP1/2 and AGAP1 inhibit the development of CDRs in NIH-3T3 cells (Jackson et al., 2000; Nie et al., 2002), whereas knockdown of ASAP1 has been shown to result in an increase in the number of CDR-containing cells (Chen et al., 2016). Furthermore, ASAP3 has been detected in CDRs (Ha et al., 2008), and in NIH-3T3 cells, ARAP1 has been established to regulate the size of CDRs via ARF1/5 (Hasegawa et al., 2012).

Certain types of cancer cells are reported to be characterized by the development of CDRs, with growth factor stimulations being shown to induce CDRs in human pancreatic cancer PANC1 cells (Orth et al., 2006), the mouse epithelial tumor Mgat5 cell line (Boscher and Nabi, 2013), and mouse melanoma 2054E cells (Kadrmas et al., 2020). Trastuzumab, a monoclonal antibody that targets human EGFR 2 (HER2), has been found to induce CDRs in the human breast cancer SK-BR-3 cell line (Bagnato et al., 2017), and similarly, in human glioblastoma LN299 cells, the growth arrest-specific 6 (GAS6) protein induces CDRs, which promotes macropinocytosis and contributes to the focal adhesion turnover in these cells (Zdzalik-Bielecka et al., 2021). Furthermore, we have recently found that growth factor treatment can induce CDRs in the human Hep3B HCC cell line, although not in other HCC cell lines, such as HepG2 and Huh7, or the LO2 hepatocyte cell line (Sun et al., 2022). Given that macropinocytosis functions in the uptake of extracellular solutes, it has been proposed that heightened nutrient uptake via macropinocytosis in cancer cells triggers sufficient cell growth resulting in tumor development (Yoshida et al., 2018a; Swanson and Yoshida, 2019; Qiu et al., 2022; Puccini et al., 2022). On the basis of these findings, we hypothesized that the development of CDRs in cancer cells is associated with abnormally high levels of macropinocytosis, thereby contributing to the uptake of excess amounts of nutrients and thus promoting rampant tumor growth (Sun et al., 2022).

Interestingly, the ARF-GAP ARAP1, which is involved in CDR formation, has also been suggested to play a role in the development of cancer. ARAP1 antisense RNA1 (ARAP1-AS1) has been found to enhance the levels of ARAP1 mRNA (Yang et al., 2019), and increases in ARAP1-AS1 expression have been reported in human gastric cancer (Jiang et al., 2020). Moreover, a low level of ARAP1-AS1 expression has been demonstrated to be associated with an inhibition of lung cancer proliferation (Tao et al., 2020), and silencing ARAP1-AS1 has the effect of suppressing the proliferation of breast cancer cells (Lu et al., 2020) and HCC cells (Zhuang Qiuyu, 2021). Although several studies have proposed the molecular mechanisms by which ARAP1-AS1 modulates tumor development such as in bladder cancer (Teng et al., 2019; Zhang et al., 2022), renal cell carcinoma (Zhong and Zhong, 2021); and ovarian cancer (Li et al., 2021), the roles of ARAP1 and ARAP1-AS1 (as the antisense RNA of ARPA1) in cancer development have yet to be sufficiently determined. It could be hypothesized that ARAP1 is involved in cancer development via CDR formation.

On the basis of the findings of the aforementioned research, in this study, we investigated the role of CDRs in the context of the cancer development by focusing on the cellular function of ARAP1 in Hep3B cells. Confocal microscopy revealed that ARAP1 is expressed in the CDRs of Hep3B cells, and we observed that the size of CDRs, extracellular solute uptake efficiency, and rate of cell growth were all reduced in cells in which ARAP1 had been knocked down. Invasion and migration assays to test the malignant potentials of the cells showed that depletion of ARAP1 attenuated the efficiency. Interestingly, confocal microscopy revealed that ARAP1 was uniquely expressed in the mitochondria and that the pattern of mitochondrial expression was altered in cells in which ARAP1 had been knocked down, thereby providing evidence for an interaction between the mitochondria and CDR formation. Furthermore, database analysis indicated that there is a strong correlation between cell survival and the expression of ARPA1 mRNA. In addition, on the basis of human tissue staining, we established that ARAP1 expression in tumor cells was higher than that in the cells of non-tumor control tissue. Collectively, our findings thus provide convincing evidence to indicate that an abnormal expression of ARAP1 in Hep3B cells induces the formation of CDRs, leading to heightened nutrient uptake that in turn triggers the development of cancer.

## RESULTS

### The GTPase-activating protein ARAP1, although not the target molecule ARF1, is located in the CDRs of Hep3B cells, and depletion of ARAP1 alters CDR size

To characterize the role of CDRs in the activity of Hep3B cells, we generated knockout (KO) cells in which the molecular mechanisms underlying CDR formation were attenuated. In this regard, we decided to generate ARAP1 KO cells, as it has previously been demonstrated that ARAP1 regulates the size of CDRs in NIH-3T3 cells (Hasegawa et al., 2012). Initially, we sought to determine whether, as in NIH-3T3 cells; ARAP1 and the target protein ARF1 are located in the growth factor (GF)-induced CDRs of Hep3B cells. Based on confocal microscopy observations, we accordingly established that ARAP1, although not ARF1, is expressed in the CDRs induced by EGF and insulin (**Fig. 1A and B**). It has previously been shown that the small GTPase ARF6 and the upstream signaling molecule NUMB1 are expressed in CDRs (Zobel et al., 2018), and in the present study, we also observed that these molecules are recruited to the CDRs in Hep3B cells (**Supplementary Fig. 1A and B**). Having initially determined these distributions, we established two different ARAP1 KO cell lines, namely, KO1 and KO2, with western blot analysis and immunofluorescence (IF) staining confirming that the expression of ARAP1 had been successfully depleted in these two lines (**Fig. 1C and Supplementary Fig. 1C**). These IF images also were useful for verifying the efficacy of the antibodies used for cell staining (**Supplementary Fig. 1C**). Using these KO cell lines and the control, we performed a CDR assay, which revealed that the size of CDRs was reduced in the KO cells (**Fig. 1D and E**). Next, we compared the size of CDRs in the control and the KO cells using the average size of CDRs in the control as 1.0. Interestingly, having defined a CDR with a size greater than 2.0 as a “large-sized” CDR and compared the individual images, we found that whereas large-sized CDRs were observed in the control cells, we were unable to detect any CDRs of this size in the KO cells (**Fig. 1E and Supplementary Fig. 1D**). These findings would thus tend to indicate that ARAP1 is associated with a molecular mechanism that contributes to determining the size CDRs. In a previous study, we showed that CDRs regulate GF-induced AKT phosphorylation (pAKT) in Hep3B cells (Sun et al., 2022), and in the present study, our western blot analysis revealed that levels of pAKT in the KO cells were lower than those detected in the control cells (**Fig. 1C**), albeit non-significantly so (**Supplementary Fig. 2A**). Furthermore, IF staining of pAKT and AKT revealed that whereas the ratio of pAKT/AKT in the CDRs of KO1 cells was less than that for control cells, this difference was not detected in the KO2 cells (**Supplementary Fig. 2B and C**). Based on these observations, we concluded that although the depletion of ARAP1 contributes to a reduction in the size of CDRs, the difference is not sufficient to influence CDR-dependent pAKT.

**Figure 1.**
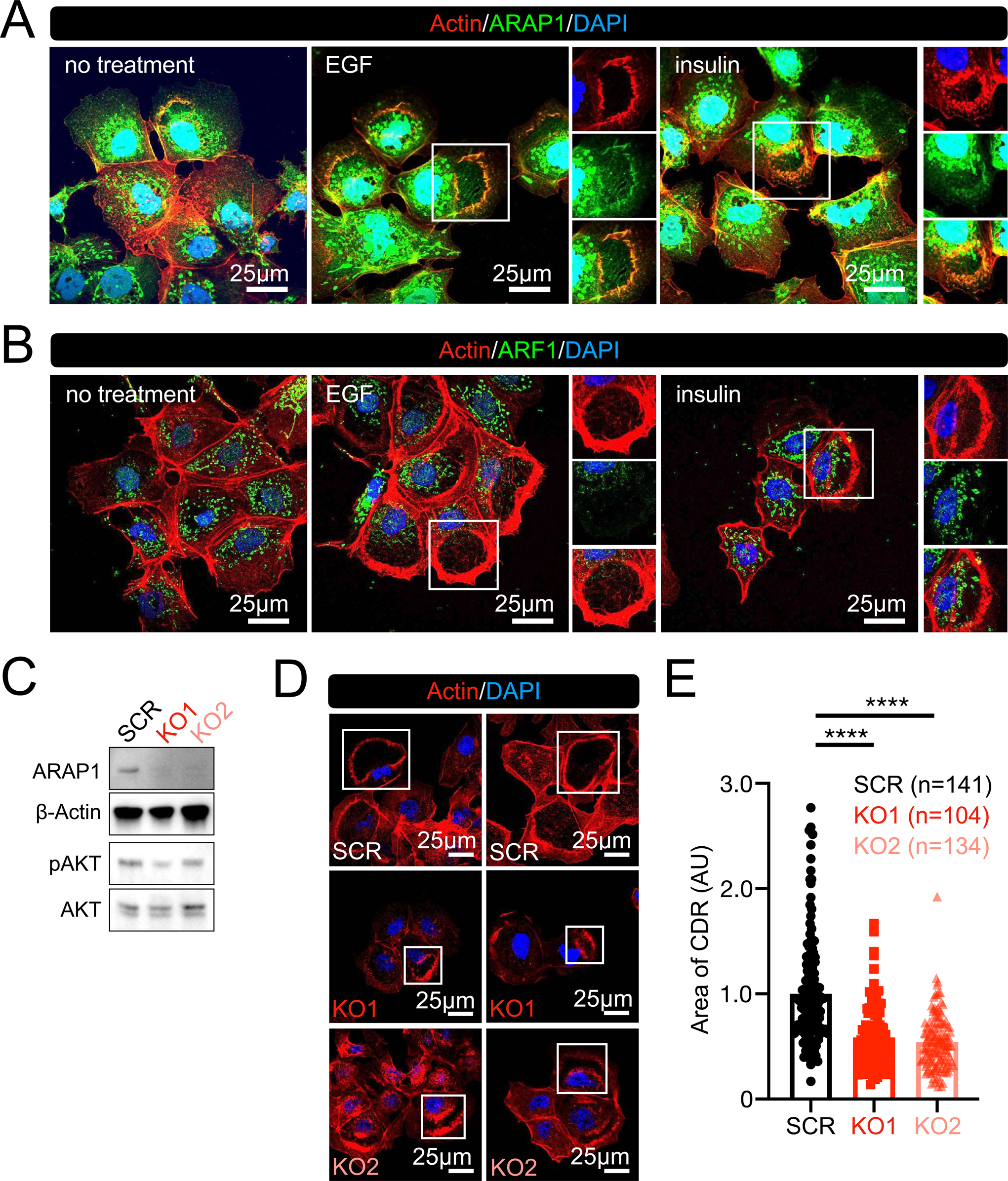
ARAP1 modulates the size of circular dorsal ruffles (CDRs) in cells of the Hep3B hepatocellular carcinoma cell line. (**A and B**) Representative confocal images of Actin/ARAP1 (**A**) and Actin/ARF1 (**B**) with/without epidermal growth factor (EGF) and insulin. Actin was used to identify CDRs. (C) Western blot analysis was performed to confirm ARAP1 knock out (KO). (**D**) Representative confocal images of CDR assays showing that the sizes of CDRs in the ARAP1 KO cells (KO1 and KO2) are smaller than those in the control cells (SCR: scramble). (**E**) Quantification analysis of the size of CDRs induced by EGF treatment in the control (SCR, n = 141), KO1 (n = 104), and KO2 (n = 134) cells from two independent experiments. The average size of CDRs in the control cells was used as the standard. \*\*\*\**p* < 0.0001 (one-way ANOVA). AU: arbitrary unit.

### Scanning electron microscopy observations revealed that CDRs comprise small-sized vertical lamellipodia, the expression pattern of which is disrupted in ARAP1 KO cells

To visualize the effects of ARAP1 depletion in the CDR formation, we performed high-resolution scanning electron microscopy (SEM). As we have reported previously (Sun et al., 2022), the SEM images obtained in the present study showed that small-size vertical lamellipodia form the spherical-like structures defined as CDRs (**Fig. 2A**). Given that our confocal observations revealed that “large-sized” CDRs were not induced in the KO cells and that the average size was reduced in these cells (**Fig. 1E and Supplementary Fig. 1D**), we also measured the area of CDRs using these SEM images. In line with expectation, we found that the average size of the CDRs in both KO1 and KO2 cells was significantly smaller that of CDRs in the control cells (**Fig. 2B**). Moreover, our examination of 33 micrographs revealed the presence of three “large-sized” CDRs in the control cells, whereas none of these structures were identified in the KO cells (**Fig. 2B and Supplementary Fig. 3A**). Interestingly, examination of high-magnification images revealed that whereas in the control cells, the small lamellipodia were regularly induced and collective formed the spherical-like structure characteristic of CDRs, the orientation of these ruffles was disrupted in the KO cells (**Fig. 2C and Supplementary Fig. 3B**). Consequently, the of SEM observations provided evidence to indicate that ARAP1 determines the size of CDRs by modulating the distribution of individual small ruffles.

**Figure 2.**
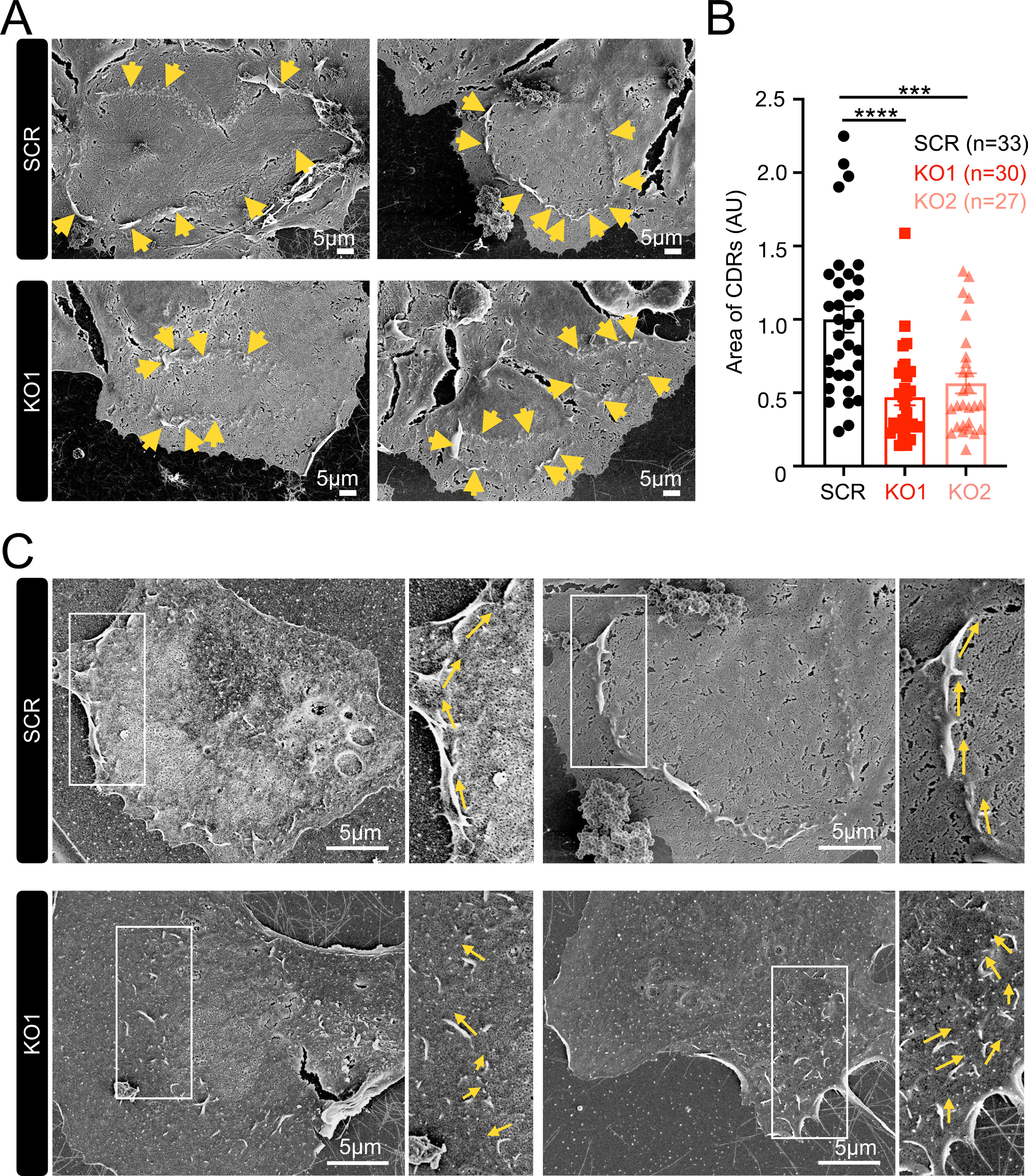
High-resolution scanning electron microscopy (SEM) observations revealed that the sizes of circular dorsal ruffles (CDRs) in knockout (KO) cells were reduced and the expression patterns of vertical lamellipodia were disrupted. (**A**) Representative high-resolution SEM images of EGF-stimulated Hep3B cells showing CDRs consisting of small vertical ruffles (arrows) in both control (SCR) and KO (KO1) cells. (**B**) Quantification of the sizes of CDRs in the control (SCR, n = 33), KO1 (n = 30), and KO2 (n = 27) cells. \*\*\*\**p* < 0.0001; \*\*\**p* < 0.001 (one-way ANOVA). AU: arbitrary unit. (**C**) Enlarged high-resolution SEM images showing details of the vertical ruffles (arrows). In contrast to the regular pattern of ruffles comprising the CDRs of the control (SCR) cells, the orientation of ruffles in the CDRs of KO (KO1) cells was disrupted. Additional images (KO2) are shown in Supplementary Fig. 3B.

### ARAP1 is mitochondrially localized and mitochondrial expression patterns are disrupted in ARAP1 KO cells

Confocal microscopy observations of ARF1 and ARAP1 signals revealed that the expression patterns of these molecules are specific, and presumably mitochondrially localized (**Fig. 1 A and B**). We thus used two well-established mitochondrial markers, TIM23 and TOM20 (Zhao and Zou, 2021; Sayyed and Mahalakshmi, 2022), to determine whether these molecules are distributed in the mitochondria. TIM23/ARAP1 double staining clearly revealed that ARAP1 was co-localized with TIM23 in Hep3B cells (**Fig. 3A**). To establish whether this expression pattern is Hep3B-specific, as a control, we performed similar analysis using cells of the Huh7 HCC cell line, in which GF does not induce CDRs (Sun et al., 2022), and accordingly failed to detect any co-localization of TIM23 and ARAP1 in these cells (**Fig. 3B and Supplementary Fig. 4A**). Given our finding of the CDR localization of ARAP1, we predicted that mitochondria might be recruited to the CDRs. Contrary to expectations, however, we observed that the expression patterns of both TIM23 and TOM20 remained virtually unaltered in response to EGF stimulation, and we detected an absence of signals within the CDRs (**Fig. 3C and D**). These observations would thus tend to indicate that following GF stimulation, ARAP1 undergoes translocation from the mitochondria to the CDRs. Moreover, our use of a negative control in this analysis enabled us to exclude the possibility that our detection of ARAP1 in the CDRs was an artifact associated with the observation method.

**Figure 3.**
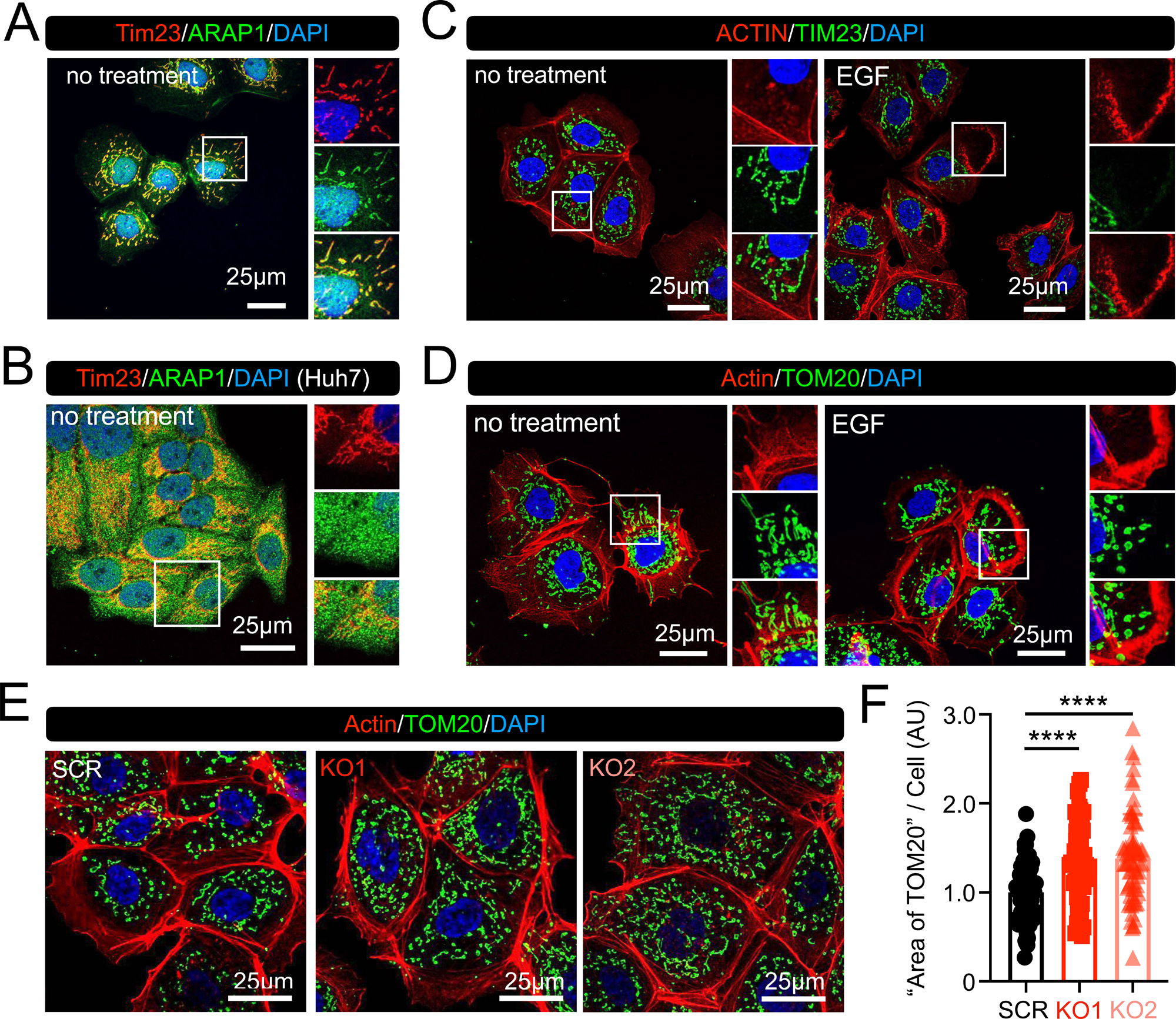
Mitochondrially localized ARAP1 modulates the pattern of mitochondrial expression pattern. (A) Representative confocal images of TIM23/ARAP1 in Hep3B cells prior to growth factor stimulation. Strong co-localization of TIM23 (red) and ARAP1 (green) was observed. (**B**) Representative confocal images of TIM23/ARAP1 in Huh7 cells without growth factor stimulation. TIM23 (red) and ARAP1 (green) were not co-localized. (**C**) Representative confocal images of Actin/TIM23 in Hep3B cells with/without EGF stimulation. TIM23 was not detected in the circular dorsal ruffles (CDRs). (**D**) Representative confocal images of Actin/TOM20 in Hep3B cells with/without EGF stimulation. TOM20 was not detected in the CDRs. (**E**) Representative confocal images of Actin/TOM20 staining in control (SCR) and knockout (KO1 and KO2) cells. The area of TOM20 expression (green) was expanded in KO cells. (**F**) Quantitative analysis of the area of TOM20 expression in different cell types. The area of TOM20 expression and sizes of cells were measured using ImageJ software and we calculated the ratio of “TOM20 expression area”/“cell area.” The results revealed that the ratios obtained for KO cells were higher than those obtained for the control cells, thereby indicating that the area of mitochondrial expression increased in response to the deletion of ARAP1. *n* = 64 (SCR), 56 (KO1), and 71 (KO2), from two independent experiments. \*\*\*\**p* < 0.0001 (one-way ANOVA). AU: arbitrary unit.

To determine whether the mitochondrial localization of ARAP1 has any influence on the pattern of mitochondrial expression in Hep3B cells, we compared the distribution of TOM20 expression in controls and KO cells. Actin was used to identify the cell area, and the ratio of TOM20 to Actin was calculated as the “area of mitochondria”/cell. Compared with the control cells, confocal microscopy revealed an expanded area of TOM20 expression in the KO cells (**Fig. 3E**). Furthermore, quantification analysis clearly showed that the ratio was slightly, albeit significantly, increased in the KO cells (**Fig. 3F**), whereas the sizes of cells remained unaltered (**Supplementary Fig. 4B**). These observations would thus tend to indicate an increase in mitochondrial area in the KO cells. By way of confirmation, we also used the mitochondrial marker TIM23, and accordingly detected an increase in the TIM23/Actin ratio in the KO cells (**Supplementary Fig. 4C-E**). Collectively, these findings indicate that ARAP1 is uniquely expressed in the mitochondria of Hep3B cells and that its expression is associated with the molecular mechanisms underlying mitochondrial expression.

### ARF1 is mitochondrially localized and inhibition of ARF1 function disrupts CDR formation

We also observed that ARF1 was co-localized with TIM23, and confirmed this co-localization in the KO cells (**Fig. 4A**). To establish whether ARF1 is involved in the mechanisms associated with mitochondrial expression, we used the ARF1 inhibitor Golgicide A (GCA) (Sáenz et al., 2009). Interestingly, we found that treatment with this inhibitor had no appreciable effects on the area of TOM20 expression (**Fig. 4 B and C**) or on the co-localization of ARF1 and TIM23 (**Supplementary Fig. 5A**). We also detected the co-localization of ARF1 and TIM23 in Huh7 cells, which would tend to indicate that ARF1 may play a role in mitochondrial expression or function (**Supplementary Fig. 5B**). Interestingly, we observed a reduction in the size of CDRs following inhibitor treatment (**Fig. 4 D and E**), and also noted the maintained CDR localization of ARAP1 in inhibitor-treated cells (**Fig. 4F**).

**Figure 4.**
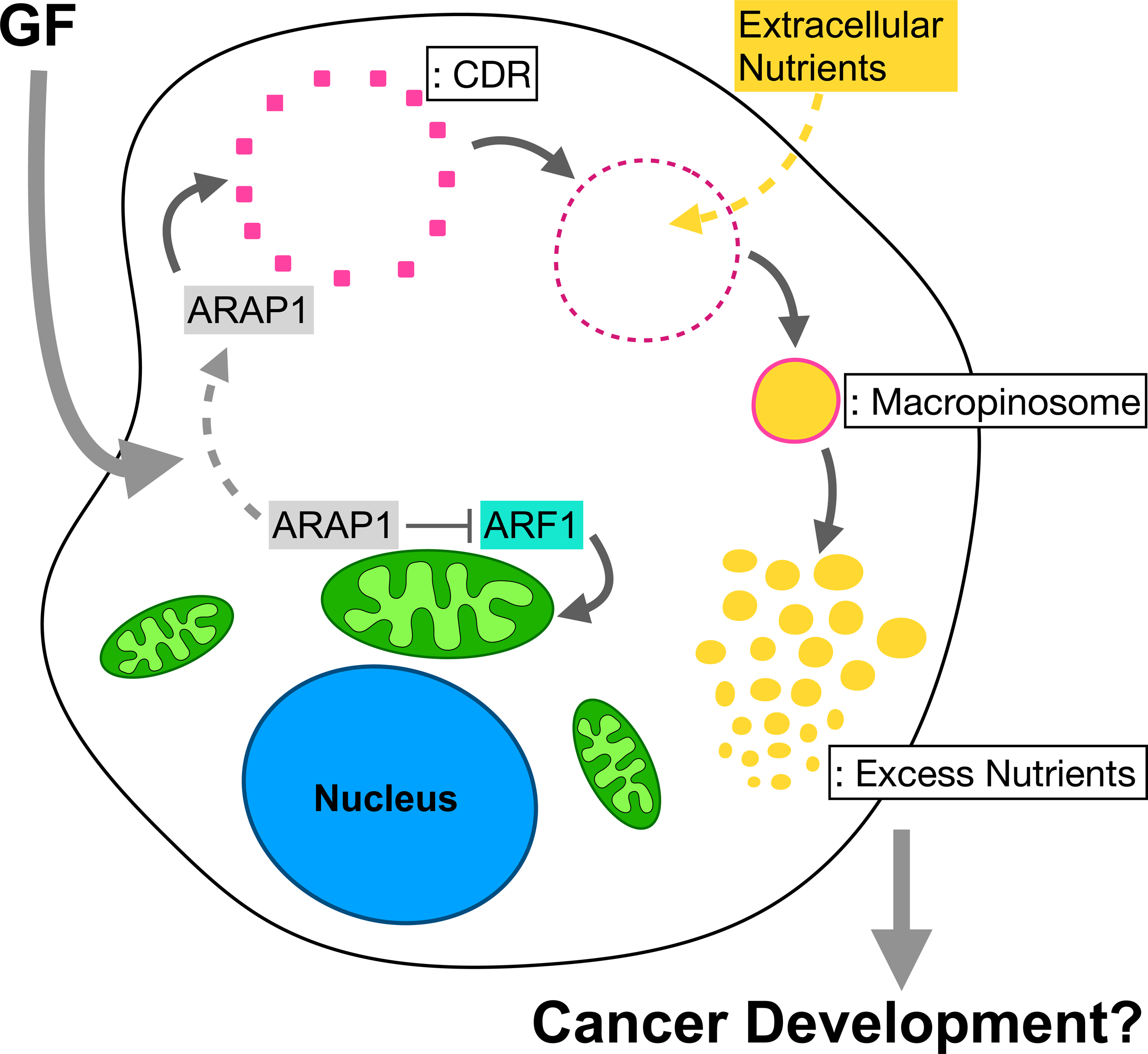
ARF1 regulates circular dorsal ruffle (CDR) size independently of ARAP1 function. (A) Representative confocal images showing the co-localization of TIM23 (red) and ARF1 (green) in both control (SCR) and ARAP1 knockout (KO1 and KO2) cells. **(B and C)** ARF1 inhibitor Golgicide A (GCA) treatment had no appreciable effects on the expression pattern of TOM20. Representative confocal images of Actin (red) and TOM20 (green) without (DMSO)/with (GCA) the ARF1 inhibitor (**B**). Statistical analysis showing that the area of TOM20 expression remained unchanged after Golgicide A treatment (**C**). *n* = 61 (DMSO) and 65 (GCA), from two-independent experiments. ns: not significant (two-tailed Student’s *t*-test). **(D and E)** GCA treatment resulted in a reduction of CDR size. Representative confocal images of EGF-induced CDRs identified by actin (red) without (DMSO)/with (GCA) the inhibitor (**D**). Statistical analysis revealed a reduction in CDR area in response to inhibitor treatment (**E**). *n* = 121 (DMSO) and 135 (GCA), from two-independent experiments. \*\*\*\**p* < 0.0001 (two-tailed Student’s *t*-test). **(F)** Representative confocal images showing that ARAP1 was localized in EGF-induced CDRs without (DMSO)/with (GCA) the inhibitor.

### Depletion of ARAP1 attenuates CDR-mediated macropinocytosis as a nutrient uptake pathway in Hep3B cells

The fluorescent dye FDx70 has previously been used as a representative extracellular solute to measure the efficiency of endocytic ingestion (Yoshida et al., 2015; Hua et al., 2023), and in the present study, we used this system to determine whether the CDRs in Hep3B cells undergo a transition to macropinosomes. Time-course experiments revealed that whereas CDRs were induced within 5 mins following EGF stimulation, they had disappeared when assessed at 15 mins, concomitant with an increase in the numbers of vesicles characterized by FDx70 signals, thereby indicating the formation of macropinosomes (**Fig. 5A and B**). These findings thus indicate that the induction CDRs is a key step in the nutrient uptake pathway of cells. By applying EIPA, which has previously been used to block macropinocytosis (West et al., 1989; Swanson and Watts, 1995; Koivusalo et al., 2010; Commisso et al., 2013), we detected a reduction in the EGP-mediated induction of macropinosomes (**Fig. 5C**), and also demonstrated that HGF treatment induces CDRs and macropinosomes (**Supplementary Fig. 6**), the formation of which was also inhibited in response to EIPA treatment (**Fig. 5D**). Subsequently, to determine whether inhibition of macropinosomes influences GF-stimulated cell growth, we performed cell proliferation assays in the presence/absence of EIPA in cell culture media containing EGF (**Fig. 5E**) or HGF (**Fig. 5F**). Compared with the control cells, we detected a reduction in the rate of cell growth in those cells exposed to EIPA. These findings thus provided evidence to indicate that by facilitating the uptake of extracellular nutrients, CDR-associated macropinocytosis plays an important role in promoting cell growth. Moreover, on the basis of our observation of smaller CDRs in ARAP1 KO cells, we hypothesized that the nutrient uptake mechanism in these cells would be attenuated. To confirm this supposition, we measured the amount of ingested extracellular solutes by comparing the intensities of integrated FDx70 detected in control and KO cells after EGF stimulation, the results of which conclusively revealed a diminished intensity of FDx70 in the KO cells (**Fig. 5 G and H**), thereby implying a disruption of the nutrient uptake mechanisms in the KO cells. Consistent with these observations, our cell growth assay revealed the rate of KO cell growth to be slower than that of the control cells (**Fig. 5 I**).

**Figure 5.**
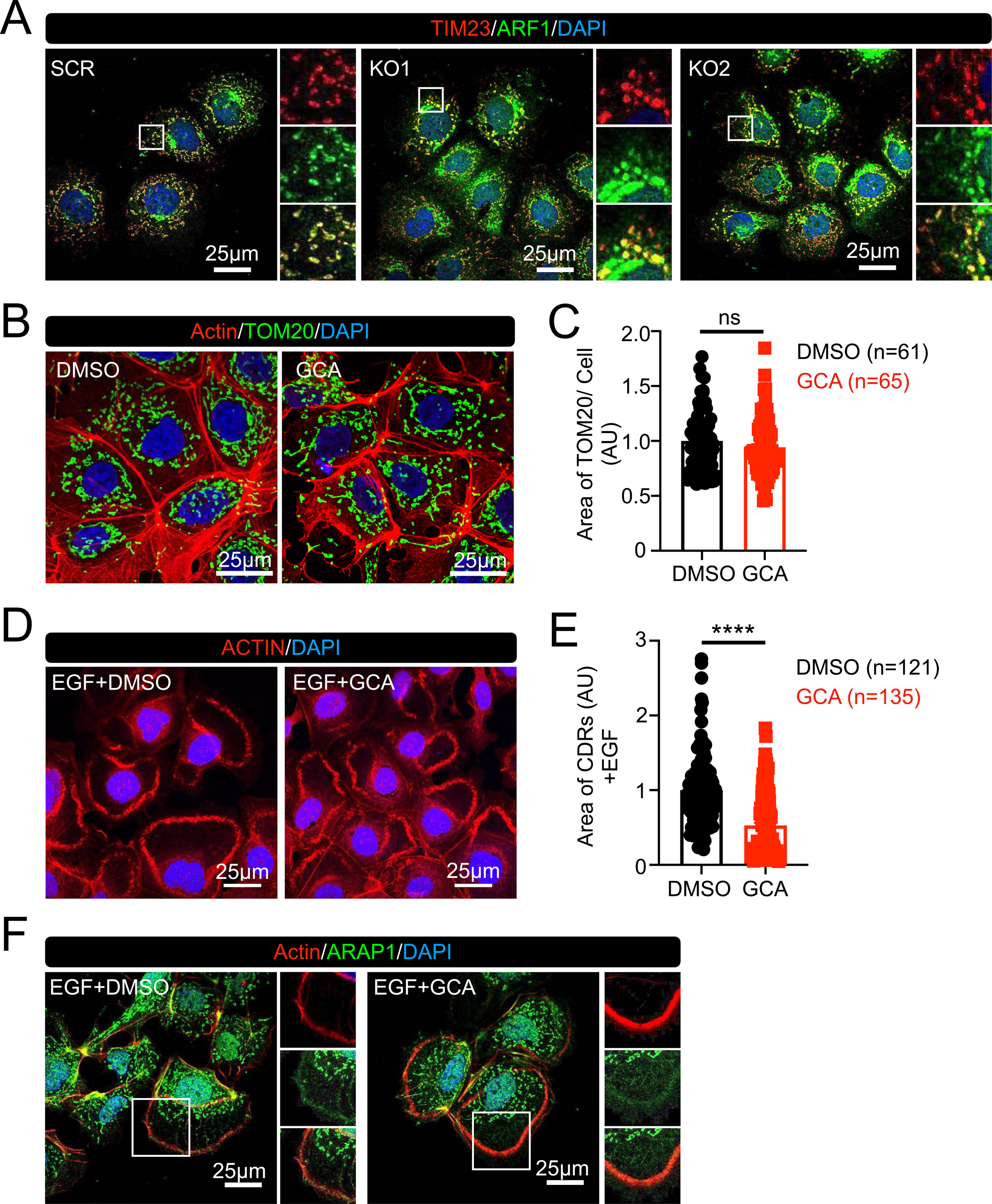
The circular dorsal ruffle (CDR)–macropinocytic process functions as a system for the uptake of extracellular nutrients. (A) Image analysis revealed CDRs (red arrows) and macropinosomes (MPs), identified using fluorescein isothiocyanate-labeled dextran with an average molecular weight of 70,000 (FDx70, green) (white arrows), in Hep3B cells following EGF stimulation. (**B**) Quantitative analysis based on three independent experiments revealed that CDRs were generated within 30 mins after EGF stimulation, with peak production occurring at 5 mins and subsequent development of macropinosomes. (**C and D**) 5-(*N*-Ethyl-*N*-isopropyl)-amiloride (EIPA) treatment inhibited the induction of MPs by EGF (**C**) and HGF (**D**). *n* = 3. \*\*\**p* < 0.001; \**p* < 0.05 (two-tailed Student’s *t*-test). (**E and F**) Results of cell counting kit-8 (CCK-8) cell proliferation assays. EIPA treatment attenuated growth factor-induced cell growth. *n* = 3. \*\*\**p* < 0.001; \**p* < 0.05 (two-tailed Student’s *t*-test). AU: arbitrary unit. (**G and H**) ARAP1 knockout (KO) cells were characterized by lower levels of nutrient uptake. Image analysis shows fewer FDx70 signals in the KOs (KO1 and KO2) cells compared with the control (SCR) cells after EGF stimulation (**G**). Quantitative analysis showing that the total intensity of FDx70 signals in the KO cells was lower than that in the control cells after EGF stimulation (**H**). FDx70 signals were measured for each cell and the values were divided by the cell area (intensity of FDx70/cell). *n* = 64 (SCR), 47 (KO1), and 58 (KO2), from two independent experiments. \*\*\*\**p* < 0.0001 (one-way ANOVA). AU: arbitrary unit. (**I**) Results of a CCK-8 assay showing that the rates of KO (KO1 and KO2) cell growth were slower than that of the control (SCR) cells. *n* = 3. \*\*\*\**p* < 0.0001; \*\**p* < 0.01; \**p* < 0.05. ns: not significant one-way ANOVA). AU: arbitrary unit.

### The involvement of ARAP1 in cancer development

Given that Hep3B is an HCC cell line, we sought to assess whether the depletion of ARAP1 would influence the malignant potentials of these cells by performing Transwell migration and invasion assays in vitro (Wang et al., 2013; Justus et al., 2014). The results clearly revealed that both the migration and invasion capacities of these cells were reduced following the knockout of ARAP1 (**Fig. 6A-D**). To further examine the potential involvement of ARAP1 in the development of cancer, we performed immunohistochemical (IHC) staining of ARAP1 using tissue samples collected from human patients. For comparative purposes, we determined the expression of ARAP1 in both tumor and control paracancerous tissues prepared from the same patients, and all staining procedures was performed under equivalent conditions. For each sample, we randomly selected five fields of view for high-magnification observations to determine IHC scores. The results revealed that the intensity of the ARAP1 signal was significantly higher in the tumor samples (**Fig. 6E and F**, **Supplementary Fig. 7**, **and Supplementary Table 1**). Moreover, database analysis revealed that there was a strong correlation between the levels of ARAP1 expression and HCC patient survival (**Supplementary Fig. 8**).

**Figure 6.**
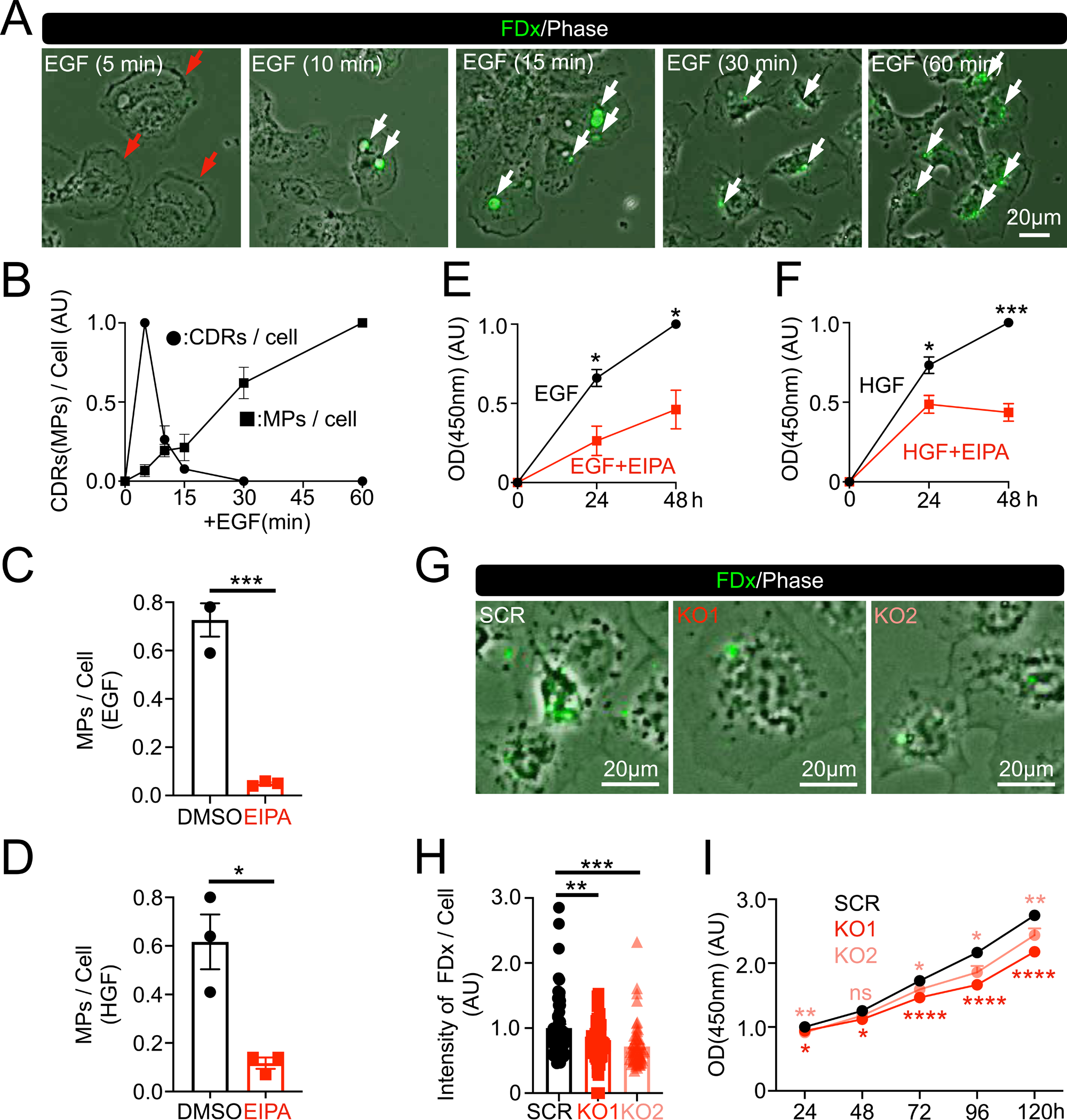
The involvement of ARAP1 in hepatocellular carcinoma tumor development. (A-D) Transwell migration (**A and B**) and invasion (**C and D**) assays in vitro showing that the deletion of ARAP1 influenced the malignant potentials of Hep3B cells. More than 10 images from two independent experiments were observed for the assays. Representative images are shown (**A and C**). The quantification results are shown as arbitrary unit (AU) (**B and D**). \*\*\*\**p*<0.0001 (one-way ANOVA). **(E and F)** Immunohistochemical (IHC) staining of ARAP1 in human hepatocellular carcinoma tissues shows that ARAP1 expression is high in tumor tissues. Tumor and paracancerous tissues (as controls) were prepared from the same patient, and five different fields of view for each tissue type were randomly selected to determine the IHC scores. Representative IHC images from two different patients **(E).** IHC staining of five patient samples were carried out independently two times (a total of 10 samples) as the 1st and 2nd sets. For quantitative analysis, the IHC scores were determined independently for each set (**F**). *n* = 5 \*\**p*<0.01 (two-tailed Student’s *t*-test). The results of the 2nd set is shown in Supplementary Fig. 7I. Other IHC images are shown in Supplementary Fig. 7A-H. Individual IHC scores are shown in Supplementary Table 1.

## DISCUSSION

In this study, we investigated the role of CDRs in the development cancer by focusing on the cellular functions of ARAP1 in Hep3B cells. We found that the CDRs in cells in which ARAP1 had been knocked out were smaller in size than those in control cells (**Fig. 1D and E and 2A and B**). Moreover, we established that the deletion of this protein attenuated the efficiency of extracellular solute uptake (**Fig. 5G and H**), rate of cell growth (**Fig. 5I**), and migration/invasion behavior (**Fig. 6A-D**) of cells. These findings thus provide evidence to indicate that by modulating the nutrient uptake pathway mediated via CDR-associated micropinocytosis, ARAP1 may play an important role in the malignant activity Hep3B cells. Consistently, our examination of stained human tissues revealed that compared with paracancerous tissues, the levels of ARAP1 protein were higher in tumor tissues obtained from the same patients (**Fig. 6E and F**, **Supplementary Fig. 7, and Supplementary Table 1**). Moreover, database analysis indicated that there was a strong correlation between the survival of HCC patients and the levels of ARAP1 mRNA expression (**Supplementary Fig. 8**). On the basis of these findings, we hypothesize that abnormal ARAP1 expression triggers an aberrant CDR/macropinocytosis response characterized by an excess uptake of nutrients, which contributes to promoting tumor development in some types of HCC (**Fig. 7**). We also observed that ARAP1 is specifically localized in the mitochondria (**Fig. 3A**). Interestingly, however, we found that the expression pattern of ARAP1 in human HCC cell line Huh7 was not mitochondria-specific (**Fig. 3B**). Given that CDRs are not produced by Huh7 cells (Sun et al., 2022), we speculate that the mitochondrial expression of ARAP1 is associated with the induction of CDR formation in Hep3B cells.

**Figure 7.**
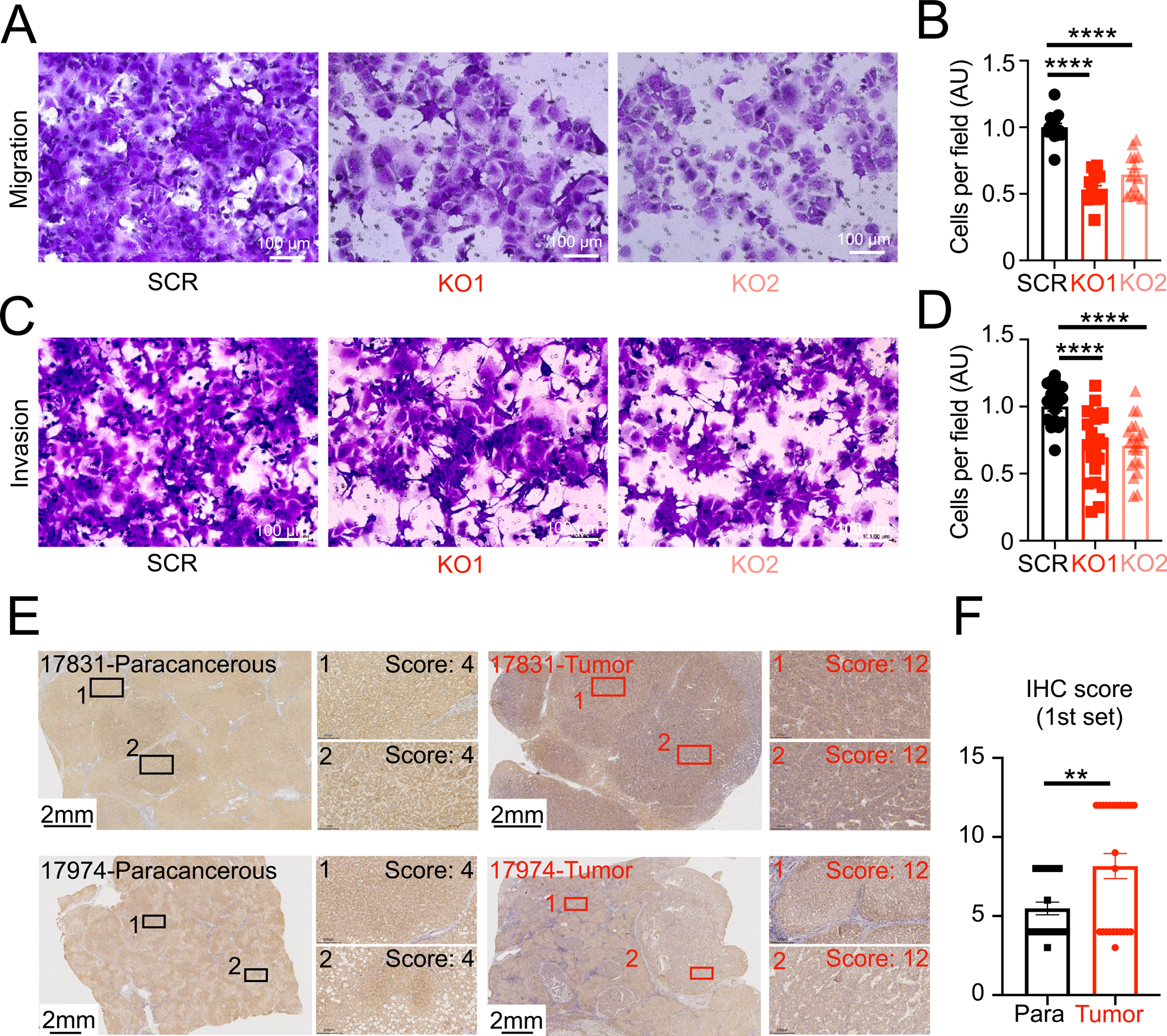
A proposed model of ARAP1 function and the physiological role of circular dorsal ruffles (CDRs) in Hep3B cells. ARAP1 is specifically expressed in the mitochondria, thereby downregulating the activity of ARF1, which modulates the pattern of mitochondrial expression. Growth factor (GF) stimulation triggers the translocation of ARAP1 from the mitochondria to the CDRs. CDRs comprise small vertical lamellipodia, the distribution of which is regulated by ARAP1. CDRs develop into macropinosomes, resulting in an excessive uptake of nutrients and thus leading to cancer development.

We found that both ARF1 and ARAP1 were at mitochondria in Hep3B cells (**Figs. 3A and 4A**). ARF1 was located at mitochondria in the ARAP1 KO cells and with ARF1 inhibitor treatment (**Fig. 4A and Supple- Fig. 5A**). Moreover, we established that whereas the deletion of ARAP1 altered the pattern of mitochondrial expression, the inhibition of ARF1 did not (**Fig. 3E and F and Fig. 4B and C**). These findings would tend to indicate that the mitochondrial expression of ARAP1 suppresses the activity of ARF1 function, thereby modulating the pattern of mitochondrial expression in Hep3B cells. Interestingly in this regard, several studies have reported that ARF1 modulates mitochondrial expression. For example, in *Caenorhabditis elegans*, activation of ARF1 by the guanine nucleotide exchange factor (GEF) GFB1 has been established to regulate the interactions between the endoplasmic reticulum (ER) and mitochondria, which results in the formation of a complex referred to as the ER–mitochondria encounter structure (ERMES) (Ackema et al., 2014; Spang, 2015), and in *Candida albicans*, the deletion of ARF1 has been found to induce the formation of ERMES (Zhang et al., 2018). The role of the GFB1–ARF1 pathway in mitochondria has also been examined in human retinal pigment epithelial RPE1 cells (Walch et al., 2018), whereas in HeLa cells, deletion of GBF1 has been demonstrated to induce the condensation of mitochondrial networks (Walch et al., 2018). In the present study, our finding that ARF1, although not ARAP1, is localized in the mitochondria of Huh7 cells (**Supplementary Fig. 5B**), indicates that the activation of mitochondrially localized ARF1 may play an important role in determining the cellular distribution of these organelles. Consequently, we hypothesize that by suppressing ARF1 activity, the expression of ARAP1 in the mitochondria disrupts the pattern of mitochondrial expression in Hep3B cells (**Fig. 7**).

On the basis of confocal microscopy observations, we established that ARAP1 is also expressed in the CDRs of Hep3B cells, which could provide evidence to indicate that in response to GF stimulation, ARAP1 is translocated from the mitochondria to the CDRs, although at present the underlying molecular mechanisms remain unclear. Interestingly in this context, it has previously been shown that mitochondrial overexpression of the ARF1-GAP ASAP1 has the effect of inhibiting the spread and migration of REF52 rat embryo fibroblasts (Liu et al., 2005). In the present study, we found that deletion of ARAP1 resulted in a reduction in the size of CDRs (**Figs. 1D and E and 2A and B**) and disruption of the expression patterns of small vertical lamellipodia in ARAP1 KO cells (**Fig. 2C and Supplementary Fig. 3B**). Thus, it can be reasoned that a downregulation of ARF1 induced by the mitochondrial expression of ARAP1 might have an influence on the cytoskeletal mechanisms associated with the formation of CDRs (**Fig. 7**). Accordingly, further studies are warranted to elucidate the molecular mechanisms and cellular function of mitochondrially expressed ARAP1 in Hep3B cells.

We also established that inhibition of ARF1 similarly resulted in a reduction in the size of CDRs, (**Fig. 4D-E**), even though we were unable to detect this protein in the CDRs (**Fig. 1B**). Given that ARAP1 was found to be localized in the CDRs, even after ARF inhibitor treatment (**Fig. 4F**), we speculate that ARF1 and ARAP1 contribute independently to the regulation of CDR formation. In previous studies that have examined the roles of ARF1 and other ARFs in CDR formation, Hasegawa et al. observed the localization of GFP-ARF1 in the PDGF-induced CDRs of NIH-3T3 cells (Hasegawa et al., 2012), and also found that overexpression of the dominant-negative forms of ARF1 and ARF5 induced larger-sized CDRs. Furthermore, it has been demonstrated that ARF6 regulates CDRs as a downstream signaling molecule of NUMB-EFA6B pathway (Zobel et al., 2018). EFA6B is one of the ARF6 GEFs (Derrien et al., 2002), and these authors revealed that EF6B is regulated by the tumor suppressor NUMB, the deletion of which (presumably leading to the deactivation of ARF6) was associated with an increase in the number of CDRs induced by HGF in HeLa cells, as well as that induced by PDGF in MEFs. It has also been shown that overexpression of the dominant-active form of ARF6 (Q67L) is associated with a reduction in the numbers of PDGF-induced CDRs in NIH-3T3 cells (Hasegawa et al., 2012). In the present study, we similarly observed that both NUMB and ARF6 are CDR localized in Hep3B cells (**Supplementary Fig. 1A and B**). Although the precise mechanisms have yet to be established, our findings provide evidence to indicate that ARFs 1, 5, and 6 coordinately regulate CDR development via GAPs and GEFs. Further studies using the same cell types and ligand sets should be conducted to elucidate the roles of each of these molecules in CDR formation.

To summarize, in this study, we characterized the roles of ARAP1 with respect to the development of circular dorsal ruffles in Hep3B cells. Our findings indicate that ARAP1 regulates both the pattern of mitochondrial expression and CDR formation, which leads us to speculate that there might be a mechanistic interaction between mitochondrial expression and CDR formation. Scanning electron micrographs revealed that CDRs comprise small-sized vertical lamellipodia, the expression pattern of which was observed to be disrupted in response to the deletion of ARAP1. Accordingly, it is conceivable that as an initial step in the induction of CDR, mitochondrial expression might determine the distribution of individual lamellipodia. As to the cellular function of these structures, our data clearly reveal that CDRs could play an important role in the nutrient uptake of Hep3B cells in response to growth factor stimulation. Given that CDRs are not exposed on the surfaces of other types of HCC cells, such as HepG2 and Huh7, or the hepatocellular cell line LO2 (Sun et al., 2022), it can be predicted that nutrients taken up via CDRs might abnormally influence the anabolic and catabolic processes of Hep3B cells. If this is indeed the case, as CDRs have been observed in certain other types of cancer cells, it is reasonable to assume that the excess uptake of nutrients is associated with cancer development. Given the important implications of these findings, we believe that the molecular mechanisms underlying the formation of CDRs in cancer cells could serve as an effective therapeutic target.

## MATERIALS AND METHODS

### Reagents and antibodies

Recombinant human EGF (AF-100-15) and human HGF (100-39H) were purchased from Peprotech (Cranbury, USA). Recombinant human insulin (M9194) was purchased from AbMole (Houston, USA). The ARF1 inhibitor Golgicide A (HY-100540) was obtained from MedChemExpress (Monmouth Junction, USA); EIPA (1154-25-2) was obtained from Tocris (Bristol, UK); a protease inhibitor cocktail (04,693,159,001) was purchased from Roche; and rhodamine phalloidin (RM02835) was purchased from ABclonal (Wuhan, China). Anti-AKT (#9272) and pAKT (Ser473) (#4060) antibodies, used for western blot analysis, were purchased from Cell Signaling Technology (Danvers, USA); anti-ARAP1 (A10466), anti-β-Actin (AC004), and anti-NUMB (A9352) antibodies, used for western blot analysis, were purchased from ABclonal; anti-ARF1 (10790-1-AP), anti-ARF6 (20225-1-AP), and anti-TOM20 (11802-1-AP) antibodies, used for immunofluorescence staining, were obtained from Proteintech (Wuhan, China); anti-TIM23 (611222). used for immunofluorescence staining, was obtained from BD Biosciences (San Jose, USA); and anti-ARAP1 (HPA012412), used for immunohistochemistry staining, was obtained from Sigma.

### Cell culture and inhibitor treatment

Hep3B and Huh7 cells were purchased from Tongpai Biotechnology Co., Ltd. (Shanghai, China) and Hunan Fenghui Biotechnology Co., Ltd (Changsha, Hunan, China), respectively, and cultured in Dulbecco’s modified Eagle’s medium (DMEM: C12430500BT; Gibco) supplemented with 10% fetal bovine serum (FBS: FS301-02; TransGen Biotech, Beijing, China), penicillin (B25911; Shanghai Yuanye Bio-Technology Co., Ltd), and streptomycin (A610494-0050; Sangon Biotech, Shanghai, China). To prevent mycoplasma contamination, cells were treated with prophylactic plasmocin (ant-mpp; InvivoGen, San Diego, USA), according to the manufacturer’s instructions. For inhibitor treatments, cells were pre-treated for 60 min with Golgicide A (10 μM).

### Establishment of a stable ARAP1-deficient Hep3b cell line

CRISPR-based knockout cell lines were generated using the lentiCRISPRv2 vector obtained from Addgene, which expresses a single guide RNA, Cas9 protein, and puromycin resistance gene. The ARAP1 single guide RNAs were designed, synthesized, and cloned into the lentiCRISPRv2 vector. Two different single guide RNAs targeting the human ARAP1 were designed [sgARAP1 1^#^: 5′-CACCGAATAGCTGCGCCACACCCCA-3′ (forward) and 5′-AAAC TGGGGTGTGGCGCAGCTATTC-3′ (reverse); and sgARAP1 2^#^: 5′-CACCGAGCCAGAGTGATGACCAAGA-3′ (forward) 5′-AAAC TCTTGGTCATCACTCTGGCTC-3′ (reverse)]. An empty lentiCRISPRv2 plasmid was used as the control vector. 293T packaging cells from Hunan Fenghui Biotechnology Co., Ltd (Changsha, Hunan, China) were seeded in 10-cm culture dishes and subsequently co-transfected with 4 μg sgRNA plasmid or empty lentiCRISPRv2 plasmid, 1 μg pMD2.G (AddGene G12259) and 3 μg psPAX2 (AddGene 12260) using Lipofectamine 2000 (Invitrogen, Carlsbad, CA, USA). Following 24-or 48-h incubations, 293T cell supernatants were harvested by centrifuging for 10 min at 1000 rpm, then filtered through 0.45-μm membrane filters. The supernatants containing the virus were used to infect Hep3b cells for 6 h. After selection with puromycin (5 μg/mL) for 3 days, the knockout efficiency was assessed by western blot analysis and the confirmed KO cells were used in subsequent experiments.

### Immunofluorescence staining and confocal microscopy

Cells were cultured overnight on coverslips in low-glucose DMEM without FBS. After stimulation with growth factors, the cells were fixed in fixation buffer A (4% paraformaldehyde in phosphate-buffered saline (PBS), pH 7.4) for 20 min at room temperature and washed in TBST (20 mM Tris, 150 mM NaCl, 0.1% Tween 20, pH 7.6). For immunofluorescence (IF) staining, cells were permeabilized in 0.1% Triton X-100 in PBS for 5 min and then incubated in blocking buffer (5% bovine serum albumin in TBST) for 30 min at room temperature. For primary antibody treatment, all primary antibodies were diluted 1:50 in blocking buffer and incubated with samples overnight at 4°C. Thereafter, the samples were washed with TBST (three times for 10 min at room temperature). As a secondary antibody treatment, samples were incubated for 2 h at room temperature with anti-rabbit IgG Alexa Fluor 488 (150081; Abcam, Boston, USA) and anti-mouse IgG Alexa Fluor 546 (Invitrogen A1003) diluted to 1:500 in the blocking buffer. The cells were then incubated for 1 h at room temperature with rhodamine phalloidin diluted 1:100 in TBST containing 5% bovine serum albumin. The samples were subsequently washed three times with TBST for 10 min at room temperature and then mounted using mounting medium containing DAPI. Microscopic observations were performed using a Leica TCS SP5 confocal microscope at the Core Facility of the College of Life Sciences, Nankai University, China.

### Cell lysate preparation and western blotting

Cell lysates were prepared as previously described (Sun et al., 2022). Briefly, the cells were lysed in cold lysis buffer (40 mM HEPES pH 7.5, 120 mM NaCl, 1 mM EDTA, 10 mM pyrophosphate, 10 mM glycerophosphate, 1.5 mM Na_3_VO_4_, 0.3% CHAPS, and a mixture of protease inhibitors) for 10 min. The lysates thus obtained were centrifuged at 13,000 × *g* for 15 min at 4°C. and the resulting supernatants were mixed with 5× sodium dodecyl sulfate-polyacrylamide gel electrophoresis (SDS-PAGE) sample buffer (#E153; GenStar, Beijing, China) and boiled for 5 min. The samples were subjected to SDS-PAGE and subsequent western blotting using the indicated antibodies.

### Scanning electron microscopy

Hep3B cells were cultured on coverslips with collagen (Type I solution from rat tail, Sigma C3867) and fixed in fixation buffer B (2.5% glutaraldehyde, 0.18 M Na_2_HPO_4_, 0.019 M KH_2_PO_4_, pH 7.2) after human EGF stimulation as previously described (Sun et al., 2022). The samples were submitted to Yimingfuxing Bio (Beijing, China) for embedding, according to standard procedures. Observations were performed using a field emission scanning electron microscope (Apreo S LoVac; Thermo Fisher) at the Central Laboratory of Nankai University.

### Macropinosome assay

Cells were cultured overnight on coverslips in low-glucose DMEM without FBS. Following the additions of EGF (1 μg/mL) or HGF (10 ng/mL) and fluorescein isothiocyanate-labeled dextran with an average molecular weight of 70,000 (FDx70, 0.5 mg/mL) (D1822; Invitrogen), the cells were incubated at 37°C for 5, 10, 15, 30, and 60 min. Thereafter, the cells were fixed with 4% paraformaldehyde in PBS at room temperature for 20 min, washed three times with DPBS (B220KJ; BasalMedia, Shanghai, China) for 10 min, and mounted. For each sample, we obtained at least 10 phase-contrast and FITC-FDx70 images using a Live Cell Station (Zeiss Axio Observer Z1) at the Core Facility of the College of Life Sciences of Nankai University. Cell numbers were determined from phase contrast images and the number of induced macropinosomes was determined by counting FDx70-positive vesicles.

### Cell Counting Kit-8 cell proliferation assay

Cell Counting Kit-8 (CCK-8) cell proliferation assays were carried out as described previously (Li et al., 2019). Briefly, cells were seeded in the wells of 96-well plates (TCP010096; BioFil, Guangzhou, China) at a density of approximately 2000 cells per well (100 μL/well), with each treatment being assessed in five replicate wells. Thereafter, cell proliferation was examined daily up to 5 days, after which. 10 μL of CCK-8 solution (CK04; Dojindo Laboratories, Kumamoto, Japan) was added to the wells followed by incubation for 2 h at 37℃. The absorbance of wells was subsequently measured at 450 nm using a Multiskan FC microplate reader (Thermo Fisher Scientific).

### Transwell migration and invasion assays

To assess cell migration and invasion, we used 24-well plates containing Transwell chambers of 8 μm size (TSC020024; BioFil). For the migration assays, cells were harvested and resuspended in DMEM without FBS at a concentration of 5 × 10^5^/mL. A 100-μL aliquot of the cell suspension was seeded in the upper chambers, and 500 μL of DMEM supplemented with 10% FBS was placed in the lower chambers. Cells were incubated for 48 h, after which the medium was removed from the Transwell chambers, non-migrating cells were removed from the upper chambers using a cotton swab. The upper chambers were then washed three times with PBS, and cells were fixed with 4% paraformaldehyde in PBS. Thereafter. the cells were stained for 20 min at room temperature with 600 μL of 1% crystal violet (C805211; MACKLIN, Shanghai, China). The upper chambers were washed a further three times with PBS (each for 10 min), and the number of the cells in the upper chambers was counted using a LeicaDFC420C light microscope. For the invasion assays, the Transwell chambers were pre-treated with Matrigel (35423; Corning) diluted 1:50 with DMEM for 1 h at 37℃, after which, 1 mL of cells was placed in the upper chamber at a concentration of 2 × 10^5^/mL. The remaining procedures were carried out as described for the migration assay.

### Human tissue staining and determination of immunohistochemical scores

Paraffin sections of human liver tissues were obtained from the Tianjin First Central Hospital, Tianjin, China. The sections were initially deparaffinized in an environmentally friendly transparent dewaxing liquid (YA0031; Solarbio, Beijing, China) and sequentially re-hydrated through an ethanol series (100%-95%-80%-60%; each for 5 min). Antigen retrieval was performed by steaming slides in a sodium citrate buffer for 30 min. Thereafter, the sections were permeabilized in 0.5% Triton X-100 in PBS for 15 min and subjected to blocking using 5% normal goat serum (SL038; Solarbio) diluted in PBS for 30 min at room temperature. The slides were then incubated overnight at 4°C with ARAP1 primary antibody (HPA012412; Sigma) diluted 1:200 in primary antibody dilution buffer provided with a mouse and rabbit specific IHC detection kit (PK1006; Proteintech). The following day, the slides were washed three times with PBS and incubated for 30 min at room temperature with goat anti-rabbit/mouse horseradish peroxidase-conjugated secondary antibodies provided with the IHC detection kit. Following incubation, the slides were washed three times with PBS, and the sections were incubated with DAB solution provided with the IHC detection kit for 50 s, followed by rinsing three times with double-distilled H_2_O. Thereafter, the sections were counterstained with hematoxylin solution (G1008; Servicebio, Wuhan, China) for 4 min and sequentially hydrated through an ethanol series (60%-80%-95%-100%; each for 5 min). The preparations were then covered with a coverslip with a drop of mounting medium, and images were obtained using a slide scan system (SQS120P; Shenzhen Shengqiang Technology Co., Ltd, Shenzhen, China) at Jingzhun Technology Co., Ltd (Tianjin, China).

To determine the IHC scores, tumor and paracancerous (control) tissues derived from the same patients were stained and observed under the same conditions. For each of two independent sample set, 10 human samples (5 tumor and 5 paracancerous tissues) were stained simultaneously. IHC scores were determined as described previously (Tong et al., 2022). The intensity of the ARAP1 signal was scored according to a four-point scale: 0 (no staining), 1 (weak light yellow staining), 2 (moderate yellow-brown staining), and 3 (strong brown staining). Similarly, the percentage area of ARAP1-positive signals in each image was scored on a five-point scale: 0 (0%), 1 (1% to 25%), 2 (26% to 50%), 3 (51% to 75%), and 4 (76% to 100%). IHC scores were obtained by multiplying the signal intensity score by the signal area score (thereby giving potential values of between 0 and 12). The IHC scores obtained for each sample group were subsequently subjected to quantitative analysis.

### Quantification

To quantify the size of mitochondria in cells, original images were converted to 8-bit images using ImageJ. Actin images were used to determine the area of cells using the “polygen selection tool.” TOM20 or TIM23 images were used to generate THRESHOLD images, from which we determined the area of mitochondria. All experiments performed in this study were conducted at least twice or with at least two biological replicates. The data are expressed as the means ± SEM. For statistical analyses, we performed a one-way ANOVA (for Fig. 1E; Fig. 2B; Fig.3F; Fig. 5H and I; Fig. 6B and D; Fig. S2A, C; Fig. S4B, D, and E), two-tailed Student’s *t*-test (for Fig. 4C and E; Fig.5 C-F; Fig.6 F), and frequency distribution (for Fig. S1D; Fig. S3A). For all analyses, a p-value of less that 0.05 was considered to signify a statistical significance.

## ACKNOWLEDGEMENT

This study was supported by a Frontiers Science Center for Cell Responses Grant from Nankai University (C029205001) and the Tianjin Key Medical Discipline (Specialty) Construction Project. XS was supported by the Tianjin Graduate Students Scientific Research Innovation Project (2021YJSB087).

## AUTHOR CONTRIBUTIONS

XS and YL designed and performed the experiments, with the support from YH, LC, JW, and LD. SY conceived the study, designed the experiments, and wrote the manuscript, with the support from ZS.

## SUPPLEMENTARY FIGURE LEGENDS

**Supplementary Figure 1.** (supplemental data for Figure 1). (A and B) Representative confocal images of Actin/ARF6 (**A**) and Actin/NUMB (**B**) with/without EGF and insulin. (**C**) Representative confocal images of Actin/ARAP1 in control (SCR) and knockout (KO1 and KO2) cells confirming the deletion of ARAP1. These images also verify that the ARAP1 antibody we used in this study is suitable for immunostaining. (**C**) Distribution of circular dorsal ruffle size in control (SCR) and KO (KO1 and KO2) cells using the same image data set shown in Figure 1E.

**Supplementary Figure 2.** Depletion of ARAP1 has a slight effect on EGF-stimulated AKT phosphorylation (pAKT). (**A**) Quantitative results for pAKT/ATK ratio values obtained from nine independent experiments (as supplemental data for Figure. 1C). Control (SCR) and knockout (KO1 and KO2) cells were stimulated with EGF, and the signals were detected by western blot analysis for ratio calculations (one-way ANOVA). (**B**) Representative confocal images of pAKT/AKT staining of Hep3B cells following EGF stimulation. (**C**) Quantitative results obtained for the pAKT/AKT ratio in circular dorsal ruffles based on confocal microscopy observations. n = 7 (SCR), 6 (KO1), and 7 (KO2) cells from two independent experiments. \**p* < 0.05 (one-way ANOVA).

**Supplementary Figure 3.** (supplemental data for Figure 2). **(A)** Distribution of circular dorsal ruffle size using the same image data set used for Figure 2B. (**B**) Representative high-resolution scanning electron micrographs of knockout (KO2) cells (as supplemental data for Figure 2C). The orientation of vertical ruffles (arrows) in the KO cells was disrupted.

**Supplementary Figure 4.** (supplemental data for Figure 3) (**A**) Representative confocal images of TIM23/ARAP1 in the Huh7 hepatocellular carcinoma cell line after EGF stimulation. (**B**) Quantitative analysis showing that cell sizes were not altered in response to deletion of ARAP1 (as supplemental data for Figure 3E and F). *n* = 64 control (SCR), 56 knockout (KO1), and 71 (KO2) cells, from two independent experiments. ns: not significant (one-way ANOVA). **(C-E)** The area of TIM23 expression was expanded in the KO1 and KO2 cells compared with the SCR cells. Representative confocal images of Actin/TIM23 in Hep3B cells (**C**). The area of TIM23 expression and the size of cells were measured using ImageJ software and the ratio of “area of TIM23 expression”/“cell area” was calculated. The results revealed that the ratios for KO cells were higher than those for control cells (**D**), although the cell areas were the same **(E)**, indicating that the area of mitochondrial expression was increased in response to the depletion of ARAP1. n = 64 (SCR), 74 (KO1), and 67 (KO2), from two independent experiments. \*\**p* < 0.01; \**p* < 0.05. ns: not significant (one-way ANOVA).

**Supplementary Figure 5.** (supplemental data for Figure 4) (**A**) Representative confocal images of TIM23/ARF1 in Hep3B cells without (DMSO)/with (GCA) the ARF1 inhibitor Golgicide A. (**B**) Representative confocal image of TIM23/ARF1 in the Huh7 hepatocellular carcinoma cell line.

**Supplementary Figure 6.** (supplemental data for Figure 5A) Image analysis identified circular dorsal ruffles CDRs (red arrows) and macropinosomes (MPs) identified using fluorescein isothiocyanate-labeled dextran with an average molecular weight of 70,000 (FDx70, green) (white arrows) in Hep3B cells following HGF stimulation.

**Supplementary Figure 7.** (supplemental data for Figure 6E and F) **(A-H)** Immunohistochemical (IHC) staining of ARAP1 in human hepatocellular carcinoma tissues revealed that ARAP1 expression was higher in tumor tissues. (I) Quantitative analysis of IHC scores for the 2nd patient sample set (as supplemental data for Figure 6F). *n* = 5. \*\**p*<0.01 (two-tailed Student’s *t*-test).

**Supplementary Figure 8.** Strong correlation between the expression of ARAP1 and patient survival. Kaplan–Meier survival curves were used to estimate the overall survival of patients in the TCGA cohort (n = 182, log rank *p* = 0.0051, HR = 1.6 *p* = 0.0057). The ARAP1-high group was characterized by a poor prognosis for liver hepatocellular carcinoma. HR: hazard ratio; TCGA: The Cancer Genome Atlas. The image is modified from the original output to adjust for presentation format.

**Supplementary Table 1.** Individual immunohistochemical (IHC) scores as supplemental data for Figure 6E and F **and Supplementary Figure 7**. The table compares the pathogenesis of tumor (tumor) and paracancerous (Para) tissues from the same patients observed under equivalent conditions. Five different fields of view from each tissue were randomly selected for the determinations of IHC scores. Staining was carried out independently in duplicate.

